# Clonal dynamics of germinal center refueling by secondary immunization

**DOI:** 10.64898/2026.05.26.728008

**Authors:** Luka Mesin, Alvaro Hobbs, Jin-Jie Shen, Juhee Pae, Ariën Schiepers, Nadine Abrahamse, Hiromi Muramatsu, Ying K. Tam, Norbert Pardi, Gabriel D. Victora

## Abstract

Many vaccine regimens involve delivery of multiple doses to the same anatomical site, such that booster doses frequently encounter germinal centers (GCs) still active from prior immunization. The consequences of this “GC refueling” to B cell clonality have not been systematically investigated. Using mouse models of mRNA-LNP vaccination combined with multicolor fate-mapping, longitudinal GC imaging, and immunoglobulin sequencing, we show that refueling triggers clonal burst-type expansion of GC-resident B cells, rather than recruiting local memory, resulting in marked focusing of GCs on the descendants of individual B cells. Refueling with a drifted antigen led to limited but detectable retraining of primary-cohort clones, although most variant-specific responses in this setting arose from newly recruited naïve B cells. These findings identify GC refueling as a distinct mode of vaccine response with implications for sequential immunization strategies against rapidly evolving pathogens.

## INTRODUCTION

Protective antibodies to a wide variety of pathogens can be induced by standard vaccination regimens, but generating antibodies capable of neutralizing antigenically diverse variants of rapidly evolving viruses remains a major challenge [1–3]. One approach to this problem is to increase the extent of somatic hypermutation (SHM) in the immunoglobulin genes of vaccine-induced B cell clones [4, 5], given that higher numbers of somatic mutations correlate with antibody affinity, breadth, and resistance to escape by viral drift [6–9]. This goal is most explicit in efforts to induce broadly neutralizing antibodies (bNAbs) to HIV, where the accumulation over years of dozens of often improbable somatic mutations appears to be a prerequisite for breadth [8–12].

Two strategies have been proposed to reach high mutational loads by vaccination: prolonging the lifespan of individual germinal centers (GCs)—the structures within which B cells undergo iterative cycles of SHM and affinity-based selection; and reactivating memory B cells (MBCs) to re-enter consecutive GC reactions by repeated immunization [4]. The latter has the advantage of allowing the sequential introduction of antigen variants to guide GC evolution towards recognizing conserved epitopes, a concept central to most proposed HIV vaccine regimens [2] and to the related problem of redirecting existing B cell responses towards emerging escape variants of influenza or SARS-CoV-2 [13, 14]. However, a practical obstacle to re-entry-based strategies is that, at least in mice, this process is highly inefficient, MBCs contributing only modestly to *de novo* secondary GCs upon boosting [15–18]. Even if such inefficiency applies only partially to humans [19], it may still pose a significant barrier to strategies requiring multiple rounds of MBC re-entry.

An alternative approach is to deliver antigen directly to a lymph node (LN) containing an ongoing GC reaction—a scenario we refer to as GC “refueling” [4, 20]. Refueling has the potential to extend GC lifespan while allowing GC selective pressures to be manipulated by sequential introduction of variant antigens. Recent data from us and others support the notion that delivering antigen to sites in which previous immune responses are still ongoing amplifies GC responses [16, 18, 20-24], with several studies reporting higher SARS-CoV-2 spike-specific responses after ipsilateral vs. contralateral mRNA boosting [21, 23, 24]. Importantly, given emerging evidence that human GCs persist for many months after immunization [25], second or third doses of many multi-dose vaccines are likely delivered at intervals short enough to encounter ongoing GCs [26], making refueling a common feature of standard vaccination schedules. Despite such ubiquity, however, there is little understanding of how refueling affects GC and antibody responses at the clonal level.

Here, using mouse models of SARS-CoV-2 spike and influenza hemagglutinin (HA) mRNA vaccination, we show that refueling with the same antigen used for priming triggers extensive clonal burst-type expansion [27, 28] of B cells already present in the GC at the time of boosting, resulting in marked focusing of refueled GCs on the descendants of individual B cells. Refueling with a moderately divergent HA variant provides limited but measurable redirection of GCs towards the refueling antigen, most consistent with selection for pre-existing crossreactivity, while the majority of the variant-specific GC response is accounted for by *de novo* recruitment of B cells from the naïve repertoire. Nevertheless, refueling elicits a detectable crossreactive response in serum. Together, our findings clarify the clonal dynamics of the GC response to antigen refueling, with implications for the design of vaccination strategies aimed at guiding antibody evolution.

## RESULTS

### Delivery of antigen to ongoing GC reactions expands both primary-derived and naïve B cells

To determine the effects of refueling on GC clonal dynamics, we used a mouse model of immunization with messenger RNA-containing lipid nanoparticles (mRNA-LNP) encoding the SARS-CoV-2 Wuhan-Hu-1 spike (S) protein (**Fig. 1A** and **S1A**). We first primed *S1pr2*-CreERT2.*Rosa26*^LSL–tdTomato^ (S1pr2-Tomato) mice, in which GC B cells can be fate-mapped by administration of tamoxifen [29], by intramuscular injection of 3 µg of mRNA-LNP into the right quadriceps. GC B cells elicited by this immunization (the “primary cohort”) were then fate-mapped by tamoxifen gavage on days 4, 8, and 12 post-prime. At 28 days, when GCs were still ongoing in the draining (right) inguinal lymph node (LN) (**Fig. 1B** and **S1B**), we boosted mice with the same mRNA-LNP formulation either ipsilaterally, thereby refueling the ongoing primary GC, or contralaterally, targeting a naïve LN as a control.

**Figure 1.**
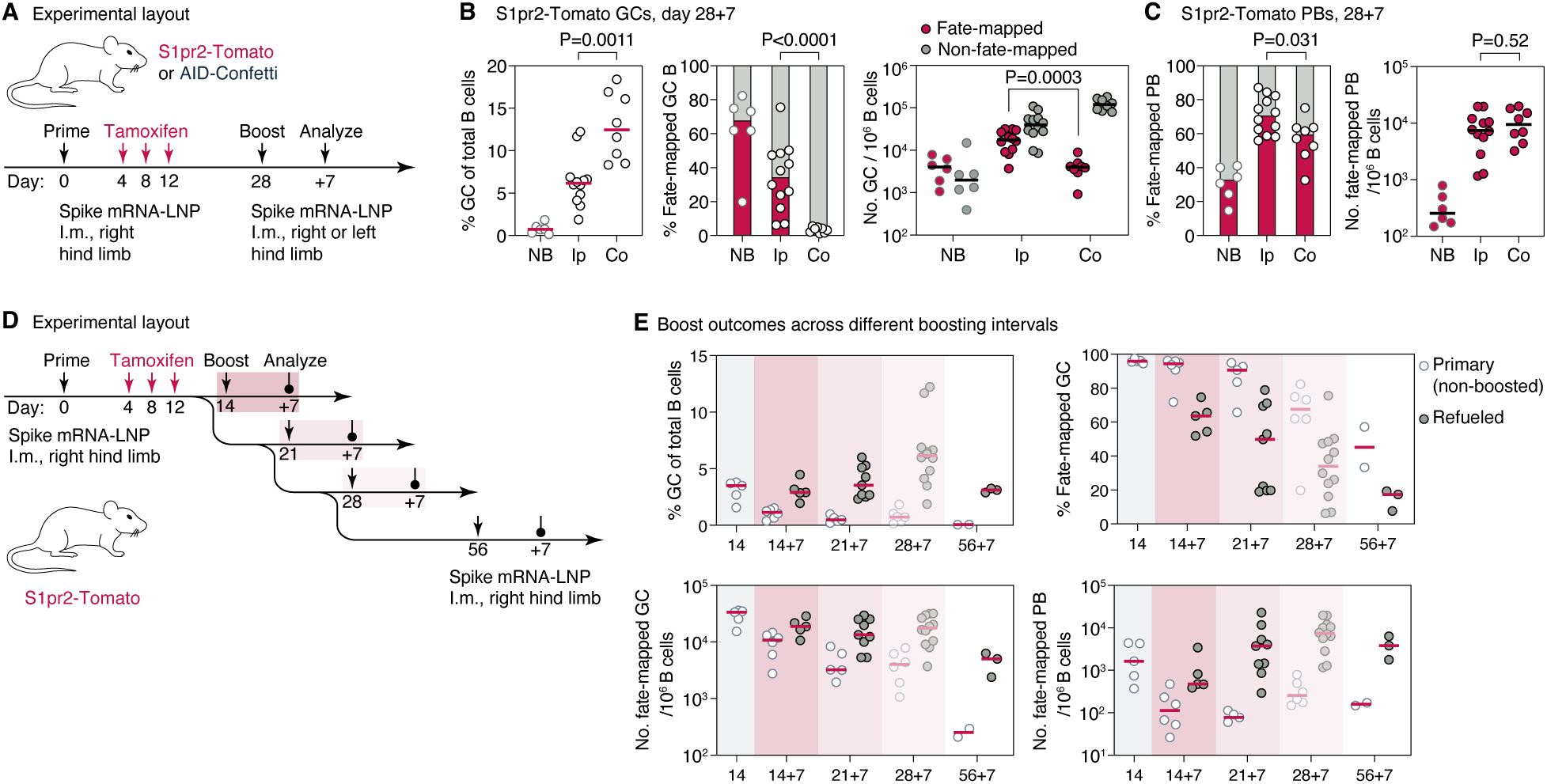
GCs refueling retains and expands primary-cohort B cell clones. **(A)** Experimental layout for (B-C). **(B)** Quantification of flow cytometry (representative plots in Fig. S1B,C) of LN samples generated as in (A), showing GC size (left), fraction of fate-mapped cells (center) and number of fate-mapped and non-fate-mapped cells (right). Each symbol represents one mouse. P-values are for Mann-Whitney U test, only relevant comparisons are shown. **(C)** As in (B), quantifying the plasmablast response. **(D)** Experimental setup for panel (E). The refueling dose varies from day 14 to 56 post-prime. Analysis is always at 7 days post-refueling. **(E)** Quantification of flow cytometry data as in (D). Each symbol represents one mouse. Primary (white) symbols are mice that did not receive a refueling dose but were analyzed at the same day as their refueled cohort. Data for the day 28+7 time point (lighter symbols) are reproduced from (B,C) for comparison. P-values are for Kruskal-Wallis test with Dunn’s post-hoc multiple comparisons test. Only comparisons with P<0.05 are shown.

Both ipsilateral refueling and contralateral boosting had pronounced effects on GC size and composition (**Fig. 1B** and **S1C**). Contralateral boosting induced large *de novo* GCs by day 7 post-boost, accounting for a median of 12% of all B cells. In agreement with our previous studies [15, 16], these GCs contained only a small fraction (3.6%) of fate-mapped, memory-derived B cells (**Fig. S1B** and **S1C**).

On the other hand, refueled GCs contained a highly variable but generally much larger fraction of fate-mapped, primary cohort B cells (median, 34%; interquartile range (IQR), 18–48%). Although this fraction was lower than that observed in non-boosted GCs at the same time point (median, 68%), the total number of both fate-mapped and non-fate-mapped GC B cells (normalized per million total B cells) increased by 4.4-fold and 20-fold, respectively, compared to non-boosted mice. Thus, refueling enlarges GCs by simultaneously expanding primary cohort B cell clones and recruiting B cells *de novo* from naïve precursors. In contrast, local plasmablast induction was remarkably similar between ipsilateral and contralateral settings: regardless of the site of boosting, the majority of plasmablasts derived from fate-mapped clones (70% and 59%, respectively) that expanded to similar numbers, suggesting that MBC behavior is similar at both sites with respect to plasmablast formation (**Fig. 1C** and **S1C**). Kinetic analysis of the ipsilateral GC and PC response showed that GC responses increased gradually from day 5 to 7 post-refueling, whereas PC numbers peaked at day 5, decreasing slightly after that (**Fig. S1D**).

To assess the effect of the timing of the refueling dose, we varied the ipsilateral boost from 14 to 56 days post-prime, assaying GCs always 7 days later (**Fig. 1D**). The fraction of fate-mapped B cells in GCs decreased progressively with later refueling (**Fig. 1E**). Between days 14 and 28, this decrease was compensated by an increase in total GC size, resulting in comparable numbers of fate-mapped GC cells per million B cells; this is consistent with proportionally greater recruitment of naïve B cells at later time points. Boosting at 56 days post-immunization resulted in smaller GCs with low fractions and numbers of fate-mapped B cells, possibly reflecting a scarcity of fate-mapped cells in the pre-boost GC due to clonal replacement [30, 31]. We conclude that the timing of refueling substantially affects GC composition and can therefore be optimized to balance primary-cohort participation with the extent of prior SHM.

To investigate the effects of GC refueling at the clonal level, we employed multicolor fate-mapping using “Brainbow” alleles [16, 32, 33] to estimate the clonal composition of various B cell compartments. We primed and boosted *Aicda*^CreERT2/+^.*Rosa26*^Confetti/Confetti^ mice (AID-Confetti) as described above, again fate-mapping GC B cells by tamoxifen treatment at days 4, 8, and 12 post-prime (**Figs. 1A** and **S2A**). As with S1pr2-Tomato mice, ipsilateral refueling resulted in a large and highly variable fraction of fate-mapped cells in boosted GCs at day 7 post-boost (median, 18%; IQR, 8.6–33%, accounting for a peak labeling efficiency of ∼60% with AID-Confetti), which substantially exceeded that observed after contralateral boosting (**Fig. S2B,C**). These differences were clear when intact LNs were visualized by multiphoton microscopy (**Fig. 2A,B**). Ipsilateral boosting produced numerous GCs densely populated with fate-mapped cells, whereas contralateral boosting yielded minimal fluorescence within B cell follicles but pronounced fluorescence in medullary regions, consistent with abundant fate-mapping of plasmablasts (**Fig. 2A,B**). Notably, GCs in ipsilaterally refueled LNs were frequently dominated by B cells of a single color (with a different color predominating in each GC; **Fig. 2A-D**), suggestive of strong clonal selection. This was evident from the high clonal dominance scores of many of the refueled GCs, 44% of which reached a score of 0.7, our typical threshold for GCs that have undergone a “clonal burst” [27], and 7 of 65 GCs consisting almost entirely of cells of a single color (NDS > 0.95), a phenotype rarely observed under conventional immunization settings [27, 28] (**Fig. 2D**). This was in contrast to the much smaller GCs found in mice prior to refueling, of which only 2 of 42 (4.8%) met the clonal burst threshold of NDS = 0.7 (**Figs. 2D** and **S2D-F**). A similar degree of post-refueling expansion was observed in mice primed and boosted with recombinant SARS-CoV-2 spike protein, ruling out an idiosyncratic effect of mRNA-LNP boosting (**Fig. S2G-K**).

**Figure 2.**
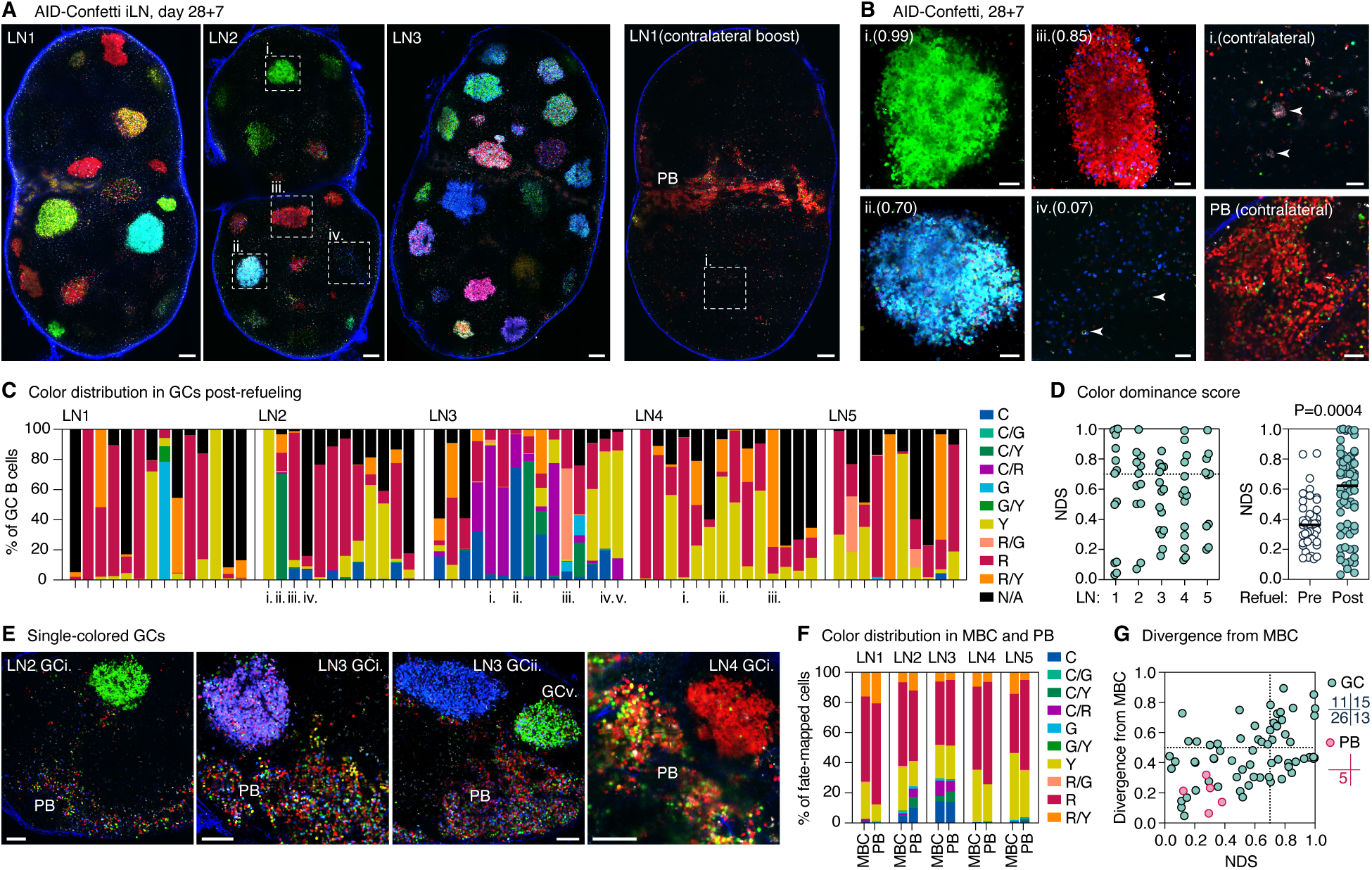
Refueling generates clonally homogeneous GCs. **(A,B)** Representative multiphoton images of cross-sections of whole LNs (A) and single GCs (B) from AID-Confetti mice at 7 days post-boosting. GCs indicated by roman numerals in (A) are shown in independently obtained magnified images in (B). Scalebars, 200 µm (A) and 50 µm (B). Numbers in parentheses are NDS. **(C)** Color distribution in individual GCs generated as in (A,B). Each bar represents one GC, colored by the fraction of cells expressing each color combination. C, CFP; G, GFP; R, RFP; Y, YFP; N/A, unlabeled cells, estimated from the density of fluorescent cells as in [27]. Roman numerals indicate GCs depicted in panels (A,B,E). **(D)** Normalized dominance score (NDS, the fraction B cells in a GC expressing the dominant color [27]) for GCs in (A-C). Each symbol is one GC. The dotted line represents a score of 0.7, the threshold for a clonal burst [27]. **(E)** Representative multiphoton images of vibratome slices of LNs in (A) (see Fig. S3), showing juxtaposition of single-colored GCs and multicolored plasmablast clusters (PB). Scalebars, 100 µm. **(F)** Color distribution of MBC and plasmablast populations in individual LNs, determined by flow cytometry. Each bar is one LN. Colors as in (C). **(G)** Color divergence (calculated as in [27]) between individual GCs and the MBC population of the same LN. Divergence between MBCs and plasmablasts (PB) from the same LNs (which quantifies the differences in (F)) is provided for comparison. Each symbol is one GC or PB sample. Quadrant lines are at 0.7 (clonal burst threshold) for NDS and at 0.5 (arbitrarily) for color divergence. The number of samples in each quadrant is given on the right.

Imaging of ∼200 µm-thick vibratome sections from these same LNs, which facilitated discrimination between GC B cells and extrafollicular/medullary plasmablasts by their position (**Fig. S3**), revealed a clear dichotomy between these two compartments: whereas GCs were largely single-colored, extrafollicular cells in peri-follicular and medullary areas (which consist primarily of MBC-derived PBs [34, 35]) displayed broad color diversity (**Fig. 2E**). Accordingly, flow cytometry of AID-Confetti mice revealed a clear convergence in colors between plasmablast and MBC populations (**Fig. 2F**). In contrast, individual GCs diverged markedly in color composition from both populations (**Fig. 2G**), often generating striking juxtapositions of single-colored GCs and multicolored extrafollicular clusters (**Fig. 2E**, and **Movie S1**). Together, these findings suggest that, in refueled LN, fate-mapped GC B cells and plasmablasts—even those positioned immediately adjacent to a GC—arise from distinct precursor pools, with plasmablasts closely mimicking the color distribution of the MBC pool, indicating a common origin.

We conclude that refueling ongoing GC responses increases GC size by both re-expanding first-cohort clones and recruiting new naïve B cells. First-cohort populations are clonally homogeneous within GCs but highly diverse within MBC and plasmablast compartments, suggesting that only rare primary-cohort cells are triggered to expand within GCs upon refueling.

### Refueling expands B cells already present in GCs prior to boosting

The unique color composition and clonal structure of refueled GCs suggested that refueling may act by triggering clonal burst-type expansion [27, 28] of B cells already present in the GC at the time the mRNA-LNP are delivered, rather than by recruiting MBCs capable of GC re-entry (and that would reside only at the ipsilateral site [18, 21], given that similar expansion is not observed contralaterally). To distinguish between these possibilities, we first varied the timing of tamoxifen administration, either treating mice during GC formation as above or delaying treatment until the week preceding the boost (**Fig. 3A**). Whereas early treatment labels GC founders and their MBC descendants, late treatment labels all cells in the GC but only a small fraction of MBCs exported between treatment and boosting (**Fig. 3B**). Refueling after late tamoxifen labeling yielded a fraction of fate-mapped GC B cells that was comparable to, if not higher than that observed with early tamoxifen (54% and 45%, respectively), consistent with expansion of fate-mapped B cells in ongoing GCs. In contrast, plasmablast labeling was substantially lower in the late tamoxifen group (29%, compared to 71% with early labeling), indicative of an MBC origin for this population (**Fig. 3C**). Thus, the kinetics of tamoxifen labeling support a GC origin for refueling-induced clonal bursts.

**Figure 3.**
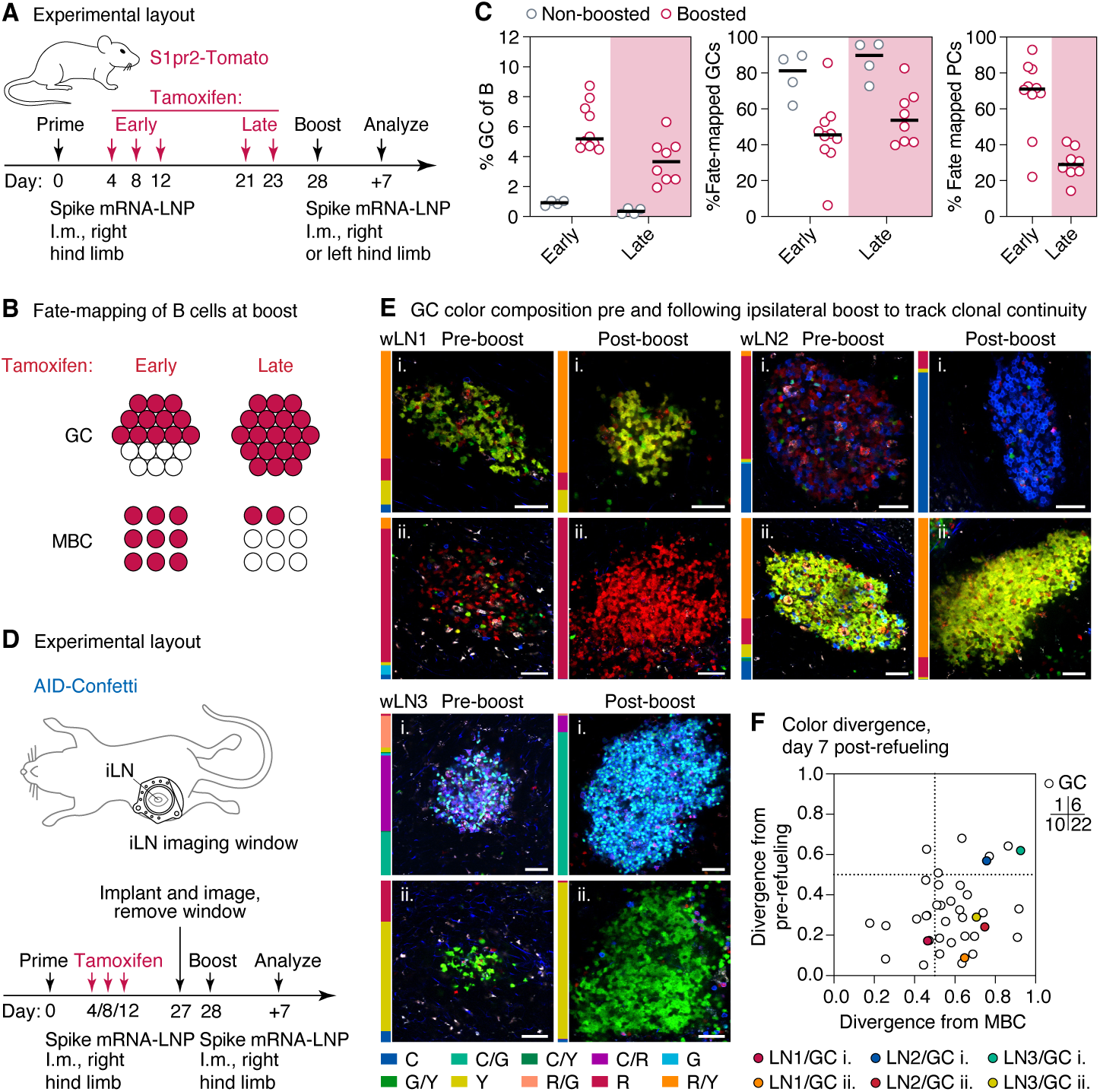
GC refueling expands clones already present in the ongoing GC. **(A)** Experimental layout for panels (B-C). **(B)** Schematic representation of the expected fate-mapping frequencies among GC and MBC populations at the time of refueling. **(C)** Quantification of flow cytometry of LN samples generated as in (A), showing GC size (left) and fraction of fate-mapped GC B cells (center) and plasmablasts (right). Flow cytometry gating as in Fig. S1A. Each symbol represents one mouse. P-values are for Mann-Whitney U test, only relevant comparisons are tested. **(D)** Experimental layout for the longitudinal imaging experiments in (E,F). GCs were imaged intravitally through implanted inguinal LN imaging windows [36], which were then removed prior to boosting. **(E)** Representative multiphoton images of selected GCs 1 day before and 7 days after refueling. Bars to the right of each image indicate the fraction of B cells of each fluorescent color in the adjacent image (color labels as in Fig. 2C). Images of all remaining pre-/post-refueling pairs are shown in Fig. S4B). Scalebars, 50 µm. **(F)** Color divergence (calculated as in [27]) between individual post-refueling GCs and either their pre-refueling images or the MBC population of the same LN. Each symbol represents one GC, with GCs shown in (E) highlighted in color. Quadrants are arbitrarily placed at 0.5 (arbitrarily) for reference. The number of samples in each quadrant is provided on the right.

To directly document the boost-driven expansion of clones already resident in the GC, we implanted mice with inguinal LN imaging windows [36, 37], which allowed us to observe the color composition of AID-Confetti GCs before and after refueling. We immunized and fate-mapped AID-Confetti mice as above and, at day 27 post-prime, implanted a titanium imaging window over the draining LN, allowing us to obtain images of GCs as they were prior to refueling. We then removed the imaging window and closed the incision. One day later, we boosted mice ipsilaterally with a second dose of spike mRNA-LNP, harvesting the LN 7 days after that to obtain post-refueling images of the same GCs (**Fig. 3D**). The GC response to refueling was similar between mice with or without implanted windows (**Fig. S4A**). Multiphoton imaging showed that pre/post-refueling GC pairs were similar in color content, and, more specifically, that post-refueling GCs in which color dominance was high invariably focused on a color that was present in that GC prior to refueling (**Figs. 3E**, **Fig. S4B**, and **Movie S2**). This included several instances of post-refueling GCs dominated by B cells expressing relatively rare color combinations (such as YFP/RFP, window (w)LN1/GC i. and wLN2/GC ii., or CFP/YFP, wLN3/GC i.) that were found frequently in the same GC at the pre-refueling time point. Quantifying divergence across all GCs showed that post-refueling GCs were much more similar (i.e., less divergent) in color distribution to their own pre-refueling state than to the MBC population in the same LN (**Fig. 3F**).

### Refueling focuses GCs on the progeny of single B cells

To understand how refueling affects GCs at the clonal level, we used single-cell sorting to isolate B cells from vibratome slices containing individual GCs (**Fig. S3)** and sequenced their *Igh* variable regions to determine clonality and somatic hypermutation (SHM) patterns [27, 28]. We first sequenced 9 GCs from the refueled LNs described in **Fig. 1**, focusing primarily (though not exclusively) on GCs with high color dominance (**Fig. 4A**). We then built phylogenetic trees from the resulting *Ighv* sequences using *gctree* software [38, 39] (**Fig. 4B** and **Fig. S5A,B**). As expected, single-colored GCs were dominated by large expanded B cell clones (**Fig. S5C**). All but one (LN4/GC ii.) of these phylogenies contained either a single or at most two nodes with the characteristic features of a clonal burst [28, 40]—multiple cells with identical sequence and a large number of close descendants, resulting from the transient halting of SHM during bursting. An extreme example was found in the large LN2 GC i. (**Fig. 2B**), in which a single clone accounted for 97% of all B cells, within which 58 of 127 sequenced cells had an identical *Ighv* gene. Extrapolating these proportions to the calculated size of that GC based on its measured volume (10^7^ µ^3^, which fits in the order of 2 × 10^4^ B cells) yielded an estimated 8 × 10^3^ cells carrying the exact same *Ighv* and 11 × 10^3^ phylogenetic descendants. Thus, refueling leads to dramatic expansion of the progeny of single B cell lineages, the largest of which can focus the entirety of the GC repertoire on a single clonal variant and its close progeny.

**Figure 4.**
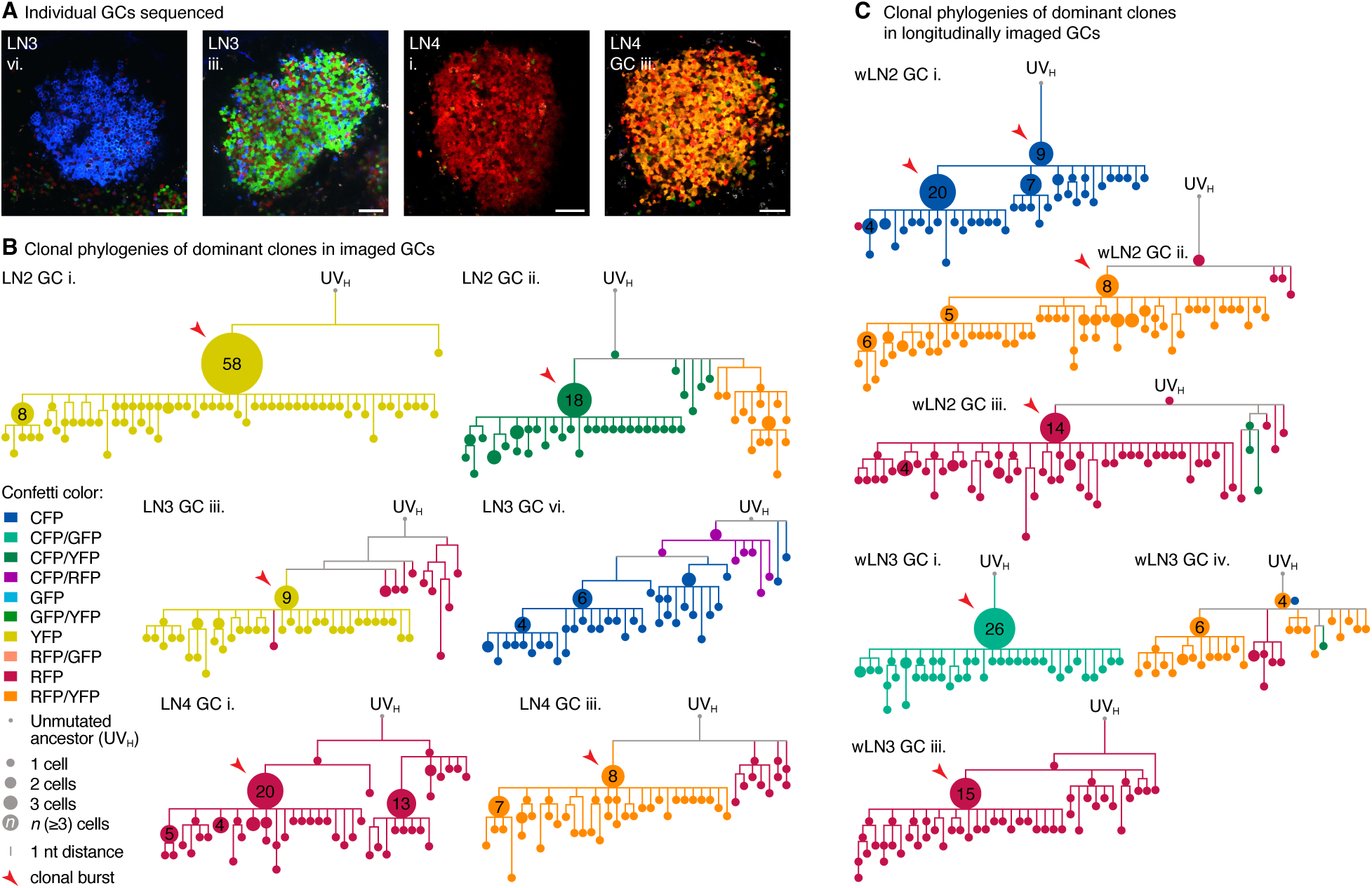
Refueling focuses GCs on the progeny of single B cells. **(A)** Multiphoton images of GCs used to build phylogenies in (B). GCs are derived from LNs analyzed in Fig. 2A-C. **(B)** Phylogenetic trees of the dominant B cell clones in the GCs depicted in (A) and in Fig. 2B. **(C)** Phylogenetic trees of the dominant B cell clones in the post-refueling time point of longitudinally-imaged GCs shown in Figs. 3E and S4B. LN and GC numbering are consistent across all figures. wLN2/GCiii. was not imaged at the pre-timepoint and is not shown as a micrograph. (B,C) Clonal bursts (defined as expanded nodes (> 1 cell) with recent expansion index (REI) > 0.25 [41]) are indicated by red arrowheads.

*S*equencing of 6 post-refueling GCs from the window experiment described in **Fig. 3D-F** confirmed these findings, in that, in every case, the phylogeny of the dominant color included at least one clonal burst-type pattern (**Fig. 4C**). Thus, color is maintained from pre- to post-refueling not by equal proliferation of all cells in the dominant color, but rather as a consequence of a “jackpot” event acting, in most cases, on a single cell (which, statistically, is more likely to carry a color frequently present in the GC prior to refueling). These dynamics are illustrated by the phylogeny of wLN3/GC i. (**Fig. 3E**), where CFP/YFP cells that were abundant but not dominant in the pre-refueling state became heavily dominant after refueling, due to a strong clonal burst initiated by a single B cell whose identical *Igh* sequence was detected 26 times among 66 sequenced members of that clone (**Fig. 4C**).

The finding that GC refueling leaves phylogenetic fingerprints in the form of large clonal bursts raised the possibility that we could detect such patterns in human GC B cells sampled before and after ipsilateral booster vaccination. We therefore analyzed previously published data [25, 42, 43] from a human clinical trial in which subjects primed with two closely-spaced doses of Pfizer-BioNTech (BNT162b2) or Moderna (mRNA-1273) SARS-CoV-2 mRNA vaccines and boosted ipsilaterally 9-10 months later with a third dose of either mRNA-1273 or mRNA-1273.213 (an experimental bivalent vaccine encoding the B.1.351 (Beta) and B.1.617.2 (Delta) spike proteins, both >98% identical to the priming variant). LNs were sampled by fine-needle aspiration (FNA) two weeks after the third dose as well as at a range of earlier time points. We identified 3 subjects with sufficient pre- and post-boost GC B cell numbers for robust clonal analysis, all of whom received mRNA-1273.213.

In each subject, spike-binding B cell clones in post-boost GCs were markedly less diverse than in pre-boost samples (**Fig. S6A**), with substantial loss from sampling of clones present at earlier time points (**Fig. S6B**) and large expansions of clones also detected prior to boosting. Phylogenetic analysis of selected expanded clones (regardless of their specificity) revealed clonal-burst-type patterns indicative of recent rapid proliferation (**Fig. S6C**) [28, 41]. Quantification of all expanded phylogenies (those accounting for ≥ 2.0% of all GC B cells in each sample) showed that clonal expansions were significantly larger in post-compared to pre-boost samples (**Fig. S6D**). We conclude that the reaction of human GCs to ipsilateral boosting is qualitatively similar to the refueling response observed in mice, suggesting that the underlying dynamics may be conserved across species.

### Refueling allows limited but measurable retraining of GC B cells towards a drifted viral antigen

The abundance of primary-cohort B cells in refueled GCs allows us to directly assess the ability of this population to adapt to a variant viral antigen. To this end, we chose a moderately antigenically distant pair of influenza virus hemagglutinin (HA) variants—A/Puerto Rico/8/1934 (HA_PR8_) and A/Michigan/45/2015 (HA_MI_)—both of the H1 subtype and 81% (403/499) identical and 89% similar (455/499) at the amino acid level. We immunized mice with HA_PR8_ mRNA-LNP, fate-mapped S1pr2-Tomato mice as in **Fig. 1A**, and then refueled GCs ipsilaterally with either HA_PR8_ or HA_MI_. To allow time for retraining, GC B cells were analyzed at 14 rather than 7 days post-refueling. Refueling with both HA_PR8_ and HA_MI_ led to increased GC size compared to non-refueled animals, although heterologously refueled GCs were slightly smaller (median, 9.8% of B cells for HA_PR8_, 5.6% for HA_MI_ and 1.9% for non-refueled LNs; **Fig. 5A**). In both cases, refueled GCs were significantly larger than primary GCs induced by either immunogen (**Fig. 5A**). Moreover, both HA_PR8_-and HA_MI_-refueled GC B cells contained reasonably large fractions of fate-mapped B cells, with the same high variability in both cases (median 12% (IQR, 6.9–16%) and 25% (IQR, 2.6–36%), respectively (**Fig. 5A**). The same trends were observed when using S1pr2-Tomato mice, although fate-mapped fractions were expectedly more efficient (**Fig. S7A**). A key difference between the conditions was the extent of GC B cell binding to HA_MI_ tetramers, which was substantial in heterologous but minimal in homologous settings (**Fig. 5B**). Notably, almost all binding to HA_MI_ in heterologously-refueled GCs was found among non-fate-mapped B cells, with only rare double HA_PR8_/HA_MI_-binders among the fate-mapped population (**Fig. 5C**). Thus, even when a large population of previously matured GC B cells was allowed to re-evolve in the presence of a second HA variant, their binding to this variant was markedly inferior to that of GC B cells specifically recruited by the variant from the naïve pool.

**Figure 5.**
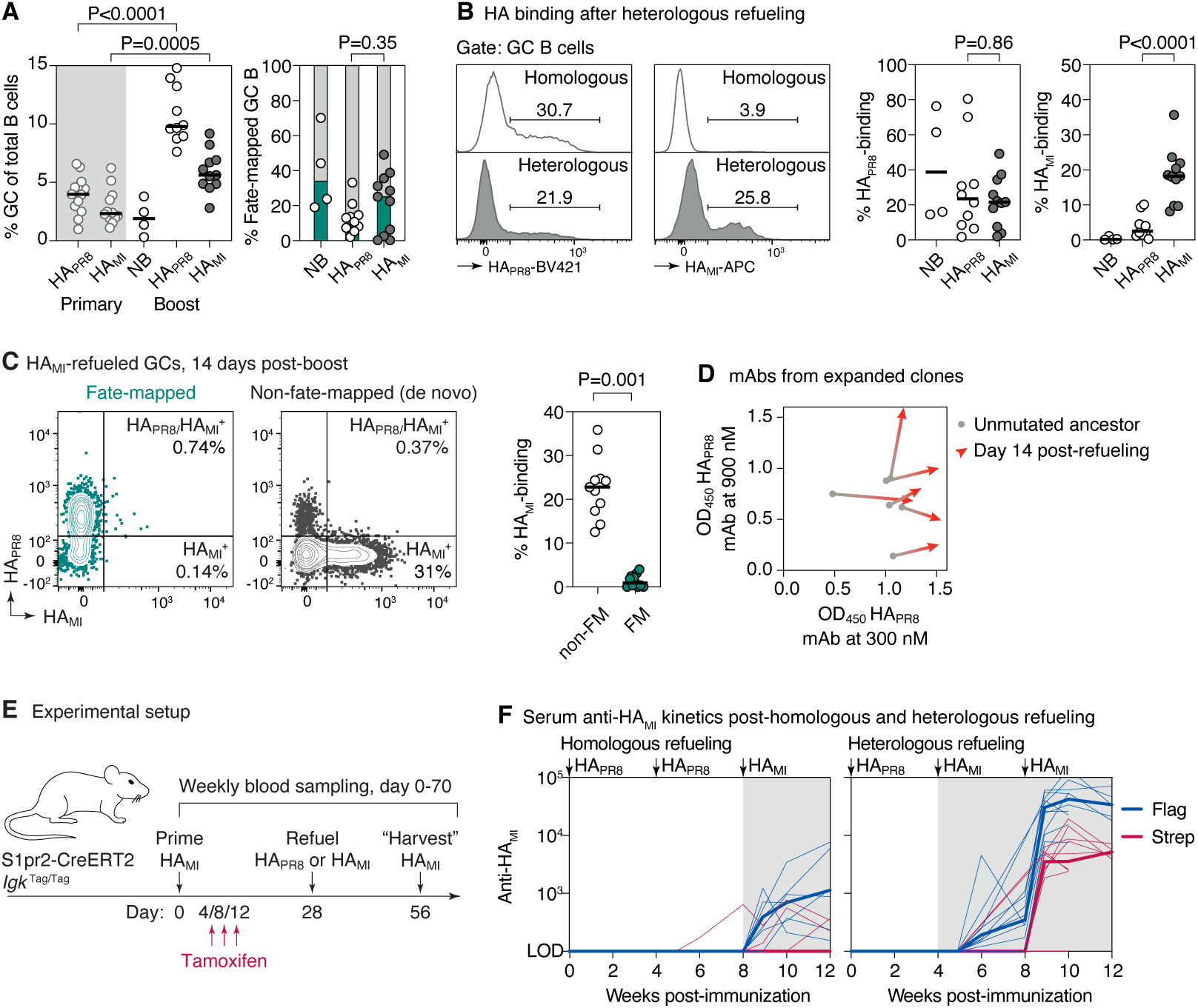
Heterologous refueling with influenza HA variants. **(A)** Quantification of flow cytometry (representative plots in Fig. S1B,C) of LN samples generated as in (Fig. 1A) using AID-Confetti mice and upon homologous or heterologous HA refueling. GC size (left) and fraction of fate-mapped cells (right) are shown. “Primary” samples were immunized once with the indicated HA variant and analyzed 14 days post-immunization. **(B)** Binding of GC B cells to the indicated HA variant by flow cytometry, quantified on the right. **(C)** Binding to the indicated HA variant of fate-mapped and non-fate-mapped GC B cells in heterologously refueled mice. (A-C) Each symbol represents one mouse. P-values are for Mann-Whitney U test, only relevant comparisons are shown. **(D)** Binding to the indicated HA variant by ELISA of mAbs derived from expanded GC B cell clones sequenced from heterologously refueled mice. Full titrations shown in Fig. S7B. **(E)** Experimental setup for (F). **(F)** Tag-specific anti-HA_MI_ serum titers in mice fate-mapped after the primary immunization (left) or at the time of refueling (right), measured by ELISA. Thin lines represent individual mice, thick lines link medians of log transformed titer values at each time point. Anti-HA_PR8_ titers are shown in Fig. S7C.

To better understand the evolution of the few primary-cohort clones that bound to HA_MI_, we isolated fate-mapped HA_PR8_/HA_MI_-binding B cells from GCs at 14 days post-refueling and produced 6 recombinant monoclonal antibodies (mAbs) from the burst points of expanded HA_MI_-refueled B cell clones, as well as their unmutated ancestors (UAs). Assaying these monoclonals against both HA variants by ELISA confirmed that they indeed bound both antigens, although binding to HA_MI_ was detectable only at higher antigen concentrations (**Fig. 5D** and **Fig. S7B**). Notably, reversion of somatic mutations revealed that all 6 mAbs were already crossreactive in their unmutated state and that, in all but one instance, SHM acted to increase binding to HA_PR8_ but not to HA_MI_. These observations are consistent with refueling selecting for B cells with pre-existing crossreactivity, rather than strictly retraining crossreactive HA_PR8_-specific B cells to bind HA_MI_.

To determine whether the limited retraining of primary-cohort clones observed in refueled GCs would be nonetheless sufficient to generate crossreactive serum antibody responses, we used our previously described “molecular fate-mapping” approach [44] to quantify HA_MI_-specific antibodies produced by first-cohort, HA_PR8_-primed B cells. In this approach, fate-mapping with an S1pr2-CreERT2 driver recombines the *Igk*^Tag^ allele, causing the Igκ constant region to switch from a Flag-tagged to a Strep-tagged state, so that antibodies produced by fate-mapped and non-fate-mapped clones can be distinguished by tag-specific ELISA. We primed S1pr2-CreERT2.*Igk*^Tag/Tag^ (K-tag) mice with HA_PR8_ mRNA-LNP, fate-mapped GC B cells at 4, 8, and 12, days post-priming, then refueled GCs at 28 days with a second dose of either homologous (HA_PR8_) or heterologous (HA_MI_) vaccine, the latter providing an opportunity for Strep-tagged GC B cells to be retrained to recognize the variant antigen. We then boosted both groups of mice with HA_MI_ in both quadriceps one month later to harvest any HA_MI_-reactive MBCs induced by the first two doses (**Fig. 5E**). Heterologous refueling generated detectable serum titers of first-cohort-derived (Strep-tagged) anti-HA_MI_ antibody that were not observed under homologous refueling, indicating some degree of productive retraining of HA_PR8_-specific B cells (**Figs. 5F** and **S7C**). Nevertheless, in agreement with our flow cytometry experiments, the bulk of anti-HA_MI_ serum antibody in the heterologous setting was Flag-tagged, indicating that it derived from the incoming naïve B cell cohort engaged *de novo* by the refueling dose. We conclude that heterologous GC refueling has the potential to partially redirect a B cell response towards reactivity to even a substantially drifted antigen. However, the naïve repertoire remains the dominant source of variant-specific GC B cells in this setting.

## DISCUSSION

We show that delivery of antigen to a LN containing an ongoing GC reaction produces effects that differ at the clonal level from those achieved by boosting a naïve LN. The most prominent effect of refueling is to trigger clonal burst-type expansion of B cells already residing in the refueled GCs, imposing a bottleneck on the diversity of B cell clones and SHM variants within this cohort. At the same time, refueling acts in a manner similar to distal boosting, engaging a sizable cohort of naïve B cells *de novo* [15, 16]. Refueling therefore leads to a superposition of old and new GC responses within the same LN.

The finding that refueling acts predominantly on GC-resident rather than memory-derived B cells is consistent with the well-documented inefficiency of MBC re-entry into *de novo* secondary GCs in mice [15–18]. Moreover, it indicates that such inefficiency also applies in settings where MBCs are present in large numbers—as they are at the ipsilateral site—suggesting that scarcity of MBCs at distal sites, or in mice in general, may not be the primary reason for the relative absence of MBCs from *de novo* secondary GCs. The clear similarity in color composition between MBC and plasmablast populations in post-refueling GCs argues that the predominant fate of refueled MBCs is to differentiate into plasmablasts locally, possibly because they are transcriptionally or epigenetically predisposed to do so [45–47]. Nevertheless, we do not exclude the possibility that local, non-migratory MBCs with greater GC re-entry capacity are present in LNs with ongoing or recently resolved GC reactions, as proposed in previous studies [18, 21]. What our data do argue is that, when antigen is delivered to LNs still harboring robust GC reactions, the contribution of MBC re-entry to the refueled GC population is minor compared to that of expanded GC-resident clones.

Our analysis of human FNA data from Alsoussi et al. [42] showed that B cell clones persisted between pre-and post-boost samples, while GCs focused on the progeny of relatively few B cells through pronounced clonal bursting. Pre-boost samples were obtained at 29 weeks post-prime for two subjects and at 15 weeks for the third, indicating that priming induced long-lived GCs in all three subjects, although we could not ascertain that GCs were indeed present at the time of boosting. Provided this is the case, the GC response to ipsilateral refueling appears to be similar in mice and humans. Another recent FNA study (Dhenni et al. [21]) explicitly compared the effects on GC formation of ipsilateral and contralateral boosting with a SARS-CoV-2 mRNA vaccine. Boosting three weeks after a contralateral primary dose resulted in negligible *de novo* GC formation 5-7 days later, in contrast to the much larger GCs observed after ipsilateral boosting both in this study and in the GC FNAs analyzed in our **Fig. S6**. This supports the interpretation that the human GCs reported by Alsoussi et al. represent refueling of GCs still present at the time of boosting, rather than *de novo* responses, although we cannot exclude that sampling in Dhenni et al. occurred too early to capture *de novo* GC formation.

Several studies have at least partially succeeded in guiding antibody responses toward broadly neutralizing epitopes of the HIV glycoprotein by sequential immunization with a series of variant antigens [22, 48–52]. Many or most of these studies involved repeated delivery of antigen to the same site and at intervals short enough to encounter ongoing GCs elicited by prior doses—conditions that, based on our findings, would favor GC refueling rather than MBC re-entry or *de novo* GC formation. Even if MBC reentry into secondary GCs is more efficient in humans than it is in mice [19], ipsilateral refueling at relatively short intervals may prove a more efficient avenue to guide antibody evolution by vaccination than strategies that depend on MBC reentry as the primary mechanism.

In addition to timing, a second variable that influences the outcome of refueling is the antigenic distance between priming and refueling proteins. Antigens too similar to the priming immunogen will predominantly expand clones targeting conserved epitopes, whereas antigens too distant will draw overwhelmingly on the naïve repertoire (at the limit, “refueling” with a completely unrelated protein generates hybrid GCs with a second specificity contributed entirely by naïve B cells [53]). Productive retraining may therefore require careful titration of antigenic distance, in a manner that mirrors the discussions of how to avoid “original antigenic sin” [15, 44, 54]. The moderately drifted HA pair used here—81% identical and 89% similar at the amino acid level, with divergence concentrated on the immunodominant head domain—appears to fall on the naïve-dominated side of this range, though detectable redirection of HA_PR8_-primed responses towards HA_MI_ was nonetheless observed at the serum level. How antigenic distance interacts with the timing of refueling to drive productive retraining is directly relevant to the design of sequential immunization strategies.

Together, our findings establish refueling as a distinct mode of GC response to vaccination, with clonal dynamics that differ qualitatively from those of either primary or *de novo* secondary GC reactions. Understanding its rules—especially how boosting interval and antigenic distance affect the clonal output of refueled GCs—may allow us to devise improved sequential immunization strategies to elicit effective antibodies against highly mutable pathogens.

## Acknowledgments

We would like to thank C. L. Ferreira for mouse work, K. Gordon and J.-P. Truman for cell sorting, V. Bahrani for help with image analysis, and the Rockefeller University Comparative Biosciences, Bioimaging, and Genomics Resource Centers and all Rockefeller University staff for their continuous support. We thank T. Okada (RIKEN/Yokohama) and T. Kurosaki (U. Osaka) for S1pr2-CreERT2 mice and C.-A. Reynaud and J.-C. Weill (U. Paris-Descartes) for *Aicda*^CreERT2^ mice. This study was funded by NIH/NIAID grants R01AI119006, R01AI173086, and R01AI180451 to G.D.V.. Work in the Victora laboratory is additionally supported by the Robertson Foundation and the Stavros Niarchos Foundation (SNF) as part of its grant to the SNF Institute for Global Infectious Disease Research at The Rockefeller University. The Pardi Laboratory was supported by NIH/NIAID grants R01AI146101 and R01AI153064.

## Author Contributions

L.M. and G.D.V. conceptualized the study. Experimental work was carried out by L.M., A.H., J.-J.S, J.P., A.S., and N.A.. N.P., H.M., and Y.K.T. generated mRNA-LNP reagents. L.M. and G.D.V. wrote the manuscript, with input from all authors.

## Use of generative AI

Claude and ChatGPT were used as aids in R coding and text editing. All generative AI output was manually verified by L.M. and G.D.V..

## MATERIALS AND METHODS

### Mice and treatments

*Rosa26*^Lox-Stop-Lox-tdTomato^ (Ai14) [55] and *Rosa26*^Confetti^ [33] mice were obtained from the Jackson Laboratories (Jax strains 007914 and 017492, respectively). *Aicda*^CreERT2^ [56] mice were provided by J.-C. Weill and C.-A. Reynaud (Université Paris-Descartes), and *S1pr2*^CreERT2^ BAC-transgenic [29] mice were provided by T. Kurosaki and T. Okada (U. Osaka, RIKEN-Yokohama). *Igk*^Tag/Tag^ mice (Jax strain 038152) [44] were generated in our laboratory. All mice were held at the immunocore clean facility at the Rockefeller University under specific pathogen-free conditions. For most experiments, 6-12-week-old male and female mice were immunized with 3 µg of mRNA-LNP intramuscularly in the quadriceps. For protein immunization, 3 µg of recombinant SARS-CoV-2 BA.1 (Omicron) spike protein (produced in-house and kindly provided by M. Nussenzweig) in alhydrogel adjuvant was also injected intramuscularly in the quadriceps. GC B cells were fate-mapped by oral gavage of 200 µl tamoxifen (Sigma) dissolved in corn oil at 50 mg/ml, on the days indicated in each experiment. Blood samples were collected by cheek puncture into microtubes prepared with clotting activator serum gel (Sarstedt).

### Lymph node imaging and sectioning

LNs were imaged as described previously [16, 27]. LNs harvested at the timepoints indicated in each experiment were first cleared of surrounding adipose tissue under a dissecting microscope then placed in PBS between two coverslips held together with vacuum grease. The tissue preparation was kept on a metal block in an ice bath throughout the duration of imaging. Multiphoton imaging was performed using an Olympus FV1000 upright microscope fitted with a 25X 1.05NA Plan water-immersion objective and a Mai-Tai DeepSee Ti-Sapphire laser (Spectraphysics) using a previously described filter set [27]. AID-Confetti fluorescence was imaged at ʎ = 930 nm. Whole-LN imaging was performed from the capsular surface (follicular cortex-facing orientation), with the exception of the PB panel in Fig. 2B, which was imaged from the hilum-facing side (medullary orientation). To image individual LN cross-sections, explanted LNs were embedded in low-melt agarose and cut into 200 μm slices using a Leica VT1000 S vibratome, as described [27]. For downstream cell sorting, LN sections were removed from agarose, further trimmed with a razor blade under a dissecting microscope to isolate fragments containing single GCs, then processed as described in the “Flow Cytometry and Cell Sorting” section, below.

### Surgical window implantation

Longitudinal window imaging of inguinal LN was performed as previously described [36, 37, 57]. Imaging windows were mounted 27 days after priming, one day prior to GC refueling. All procedures were carried out under sterile surgical conditions with mice maintained under isoflurane anesthesia. The surgical area surrounding the inguinal LN was shaved and washed with ethanol and betadine. The LN was exposed by an incision through the skin, and a custom-made titanium window fitted with a microscope coverslip was mounted as described [36]. Mice were then placed under an Olympus FV1000 multiphoton microscope on a stage specially equipped with a fixture for stable window positioning. Individual GCs were imaged in 3D as described in the previous section. After imaging, the window was removed and the skin incision was closed by suturing. Mice received subcutaneous Meloxicam ER (6 mg/kg) and were allowed to recover for 24 h before subsequent boosting with mRNA-LNPs in the quadriceps ipsilateral to the surgery site.

### Image analysis

To measure color dominance for individual AID-Confetti GCs, cells of each color combination were counted in ImageJ across 2 z-planes at least 20 µm apart. To derive the normalized dominance score (NDS) to approximate the fraction of B cells in each GC that carry the dominant color combination, the fraction of all recombined cells was first estimated in multiphoton microscopy images by calculating the density of fluorescent cells per area unit (100 µm^2^) in anatomically defined dark zone areas in which cell distribution was homogenous and dense. The density was then multiplied by the fraction of cells of the dominant color. The divergence index, which estimates the magnitude of changes in clonal distribution in GCs, was calculated as the sum of the absolute differences in percentage points between the observed proportions of each of the 10 colors in a given GC (% color * GC density) and the expected proportions in the absence of selection, as in [27]. Results were divided by 100. GC volume was measured in Imaris 10 software (Oxford Instruments) by generating a surface surrounding all fluorescent cells within a densely packed GC.

### Flow cytometry and cell sorting

LNs were processed into single-cell suspensions by maceration with a disposable micropestle (Axygen) in 100 µl of PBS supplemented with 0.5% BSA and 1mM EDTA (PBE) containing 0.5 µg Fc Block (Bio-X-Cell) in a 1.5 ml microcentrifuge tube. 100 µl of a 2X fluorescent antibody stain (see **Table S1**) was then added to the tube and cells were incubated for 30 min on ice. Cells were filtered and washed with PBE before processing for analysis or sorting on BD FACS Symphony A5, Symphony S6, or Aria II cytometer. Data were analyzed using FlowJo. For single-GC sorting, dissected fragments of vibratome slices containing individual GCs were processed and sorted as above. The remaining LN fragments not used for single-GC sorting were then combined to sort LN plasmablast and MBC samples.

### Single-cell *immunoglobulin* sequencing

Single-cell *Ig* sequencing was performed as previously described [16]. Briefly, single B cells were index-sorted into 96-well plates containing 5 μl TCL buffer (Qiagen) supplemented with 1% β-mercaptoethanol. Nucleic acids were isolated using SPRI beads, and RNA was reverse transcribed into cDNA using RT Maxima reverse transcriptase (Thermo Scientific) with oligo-dT primers. *Ig* heavy chains were amplified by PCR using a pool of V-region forward primers together with isotype-specific reverse primers. *Ig* kappa light chains were amplified separately when needed for clonality confirmation or antibody production. Primer sequences are provided in **Spreadsheet S1**. PCR amplicons were indexed with plate- and well-specific barcodes and further amplified to incorporate Illumina paired-end sequencing adaptors. Finally, amplicons were pooled, purified with SPRI beads (0.7x volume ratio), and sequenced on the Illumina Miseq platform using a 500-cycle Reagent Nano kit v2 according to the manufacturer’s instructions.

### Sequence analysis

*Ig* sequences were aligned and analyzed as previously described [16]. Phylogenetic trees were constructed using *gctree* v4.3.0 using standard settings [38, 39]. The first tree produced for each GC was used. The recent expansion index (REI), a measure of how dominated a phylogeny is by a recent clonal expansion, was calculated essentially as in [41]. Briefly, for each node in the phylogeny, the REI is the sum of the number cells on or below that node, weighted using a decay factor *τ* = 0.5, such that cells at 0, 1, 2, … nucleotide distance from the node of interest are weighted 1, 0.5, 0.25, … The total (S) is then divided by the number of sequenced cells from that GC. Only cells of the same color as the dominant node are included in the REI calculation. For human B cells, because they originate from mixtures of multiple GCs, S is divided by the number of cells in the sample, yielding a “global” REI (gREI), the values of which are thus much smaller than those of the REIs calculated for individual mouse GCs. The N75 is calculated as the number of clones accounting for 75% of cells in a sample. Chao1 is calculated as described [58]. For human LN FNA data, clonality and specificity assignments used are those given in [43].

### Monoclonal antibody production

Monoclonal antibodies were synthesized, cloned, and produced recombinantly by GenScript.

### Production of mRNA-LNP

The WH1 S mRNA-LNP were designed and encapsulated as described in [44], based on the prefusion-stabilized “2P” variant of the Wuhan-hu-1 or BA5.1 S protein sequences. HA_PR8_ and HA_MI_ mRNA-LNP were designed based on sequences provided in **Spreadsheet S2**. Coding sequences were codon-optimized, synthesized and cloned into an mRNA production plasmid (GenScript) as described in [59]. Encapsulation of the purified mRNAs in LNP was achieved using a self-assembly process where an aqueous solution of mRNAs at acidic pH is rapidly mixed with a solution of lipids dissolved in ethanol. LNP used in this study consisted of a mixture of the ionizable cationic lipid (pKa in the range of 6.0–6.5, proprietary to Acuitas Therapeutics)/phosphatidylcholine/cholesterol/PEG-lipid as described in the patent application WO 2017/004143. LNP size was measured using a Malvern Zetasizer (Malvern Panalytical, Worcestershire, UK) and encapsulation assessed using the Ribogreen assay (Life technology, Carlsbad, CA, USA).

### Recombinant antigens

HA_PR8_ was produced in house as described [16], as a cysteine-linked trimer [60] with a C-terminal foldon domain followed by an AviTag. HA_MI_ and SARS-CoV-2 RBD were purchased from SinoBiologicals (Cat# 40567-V08H1 and 40592-V08H, respectively). Antigen tetramers for flow cytometry were generated by first biotinylating antigens either site-specifically (HA_PR8_) or non-site-specifically (HA_MI_, biotinylated using the EZ-Link Sulfo NHS-LC-LC Biotin kit (ThermoFisher) at low conjugation ratio) then incubating biotinylated antigens with labeled streptavidin at 1:4 (HA:SA) molar ratio immediately prior to staining.

### ELISA

Tag-specific serum ELISAs were performed as previously (*39*). Flag and Strep ELISAs were done side by side with internal standards in each 96 or 384-well plate. Plates were coated overnight at 4 °C with 2 µg/mL recombinant SARS-CoV-2 spike or 2 µg/mL HA variants (as described in Recombinant antigens, above). Flag/Strep standard curves were added to each plate, coated with 10 µg/ml purified IgY (Exalpha Biologicals). Plates were washed with PBS + 0.05% Tween-20 (Sigma-Aldrich) and blocked for 2 h at room temperature with 2.5% BSA in PBS. Serum samples were diluted 1:100 in PBS + 0.05% Tween-20 + 0.5% BSA and serially diluted in 3-fold steps. Three-fold dilutions of a known concentration of recombinant Flag- and Strep-tagged mouse anti-IgY mAbs were added as standards.

Samples were incubated for 2 h, washed with PBS-Tween, and then incubated for 30-45 min with HRP-detection antibodies specific for each tag (rabbit anti-Flag-HRP (D6W5B) or mouse anti-Strep (Strep-tag II StrepMAB-Classic)), at dilutions pre-determined to yield identical standard curves for each tag. Samples were again washed with PBS-Tween then developed with 3,3′,5,5′-tetramethylbenzidine substrate (slow kinetic form, Sigma-Aldrich) until the reaction was stopped with 1 N HCl. Corrected Optical density (OD) values were obtained by subtracting the background absorbance at 600 nm from substrate absorbance measured at 450 nm using the Agilent BioTek Synergy HTX multimode plate reader. To normalize Flag and Strep titers, threshold OD values for the respective tagged monoclonal antibodies were first determined at fixed concentrations of either 20 or 6.67 ng/µL. Serum end-point titers were then calculated by logarithmic interpolation of the difference between the dilution readings immediately above and below the monoclonal antibody OD threshold.

For mAb ELISAs, plates were similarly coated overnight at 4 °C with 2 µg/mL of the respective HA variants. Purified mAbs were diluted to a starting concentration of 900 nM in PBS-Tween and serially diluted in 3-fold steps. Incubation, washing, HRP-detection, and development steps were identical to those described for serum ELISAs

### Data analysis

In all tests, only conditions of interest were compared, as indicated in the figure legends. Comparisons between two conditions were calculated by Mann-Whitney U test. Comparisons between more than two conditions used the Kruskall-Wallis test with Dunn’s post-test. Statistical analysis was performed using GraphPad Prism v.11. The Chao1 index was calculated in R. Flow cytometry analysis was carried out using FlowJo v.10 software. Graphs were plotted using Prism v.11 or R and edited for appearance using Adobe Illustrator 2025.

## SUPPLEMENTAL FIGURES AND LEGENDS

**Figure S1.**
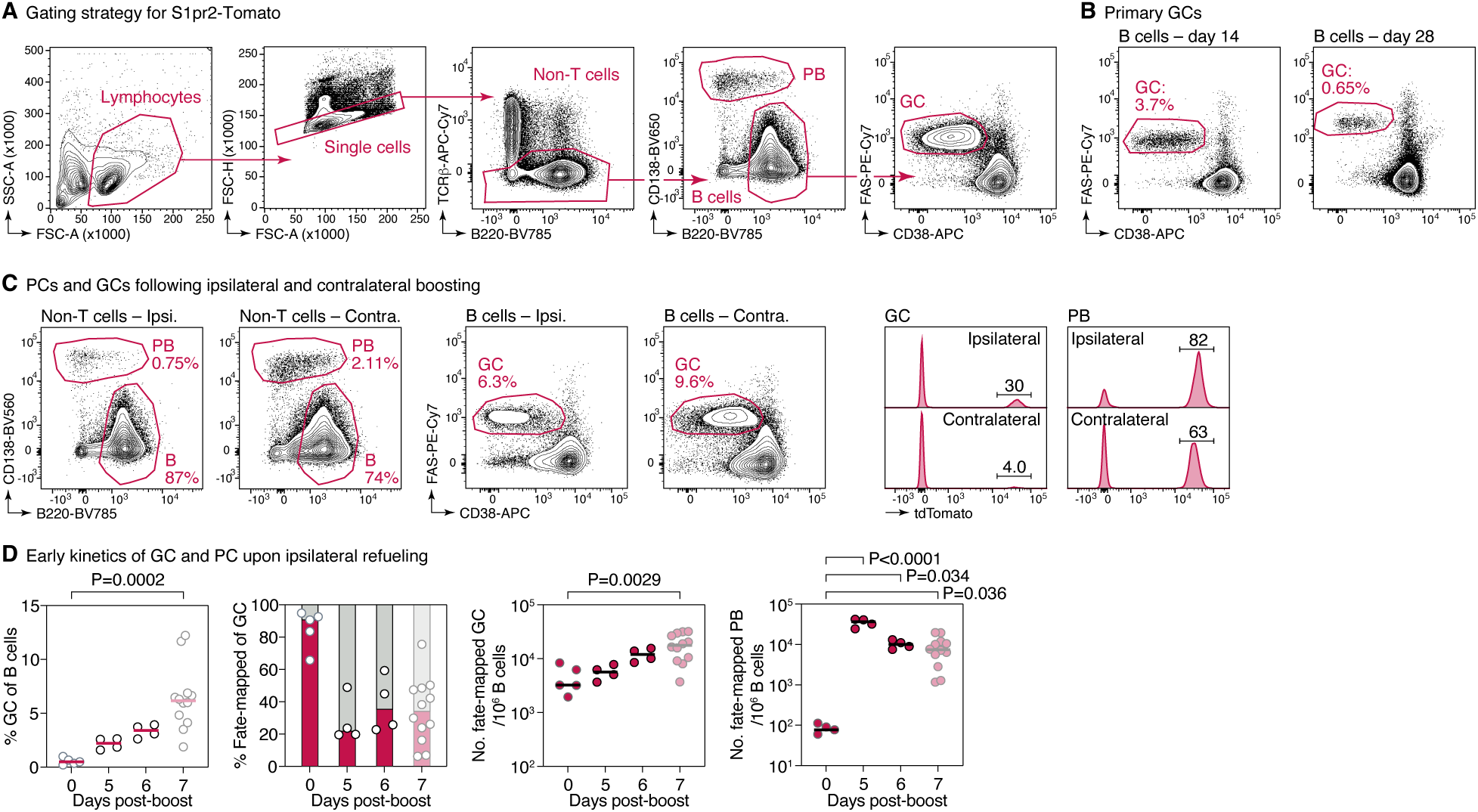
Response to refueling in mouse LNs after SARS-CoV-2 spike mRNA-LNP immunization. **(A)** General gating strategy S1pr2-Tomato fate-mapping. **(B)** Representative flow cytometry plots showing persistence of mRNA-LNP-induced GCs at the time of refueling. Gated on B cells as in (A). **(C)** Representative flow cytometry plots showing the effects of ipsilateral and contralateral on plasmablast (right) and GC B cell (center) numbers and on the fraction of fate-mapped cells within these two populations. Quantified in Fig. 1B,C. **(D)** Evolution of GC and PC expansion upon boosting. Based on flow cytometry data as in (A-C). Each symbol represents one mouse. Data for the day 7 time point (lighter symbols) are reproduced from Fig. 1B,C for comparison.

**Figure S2.**
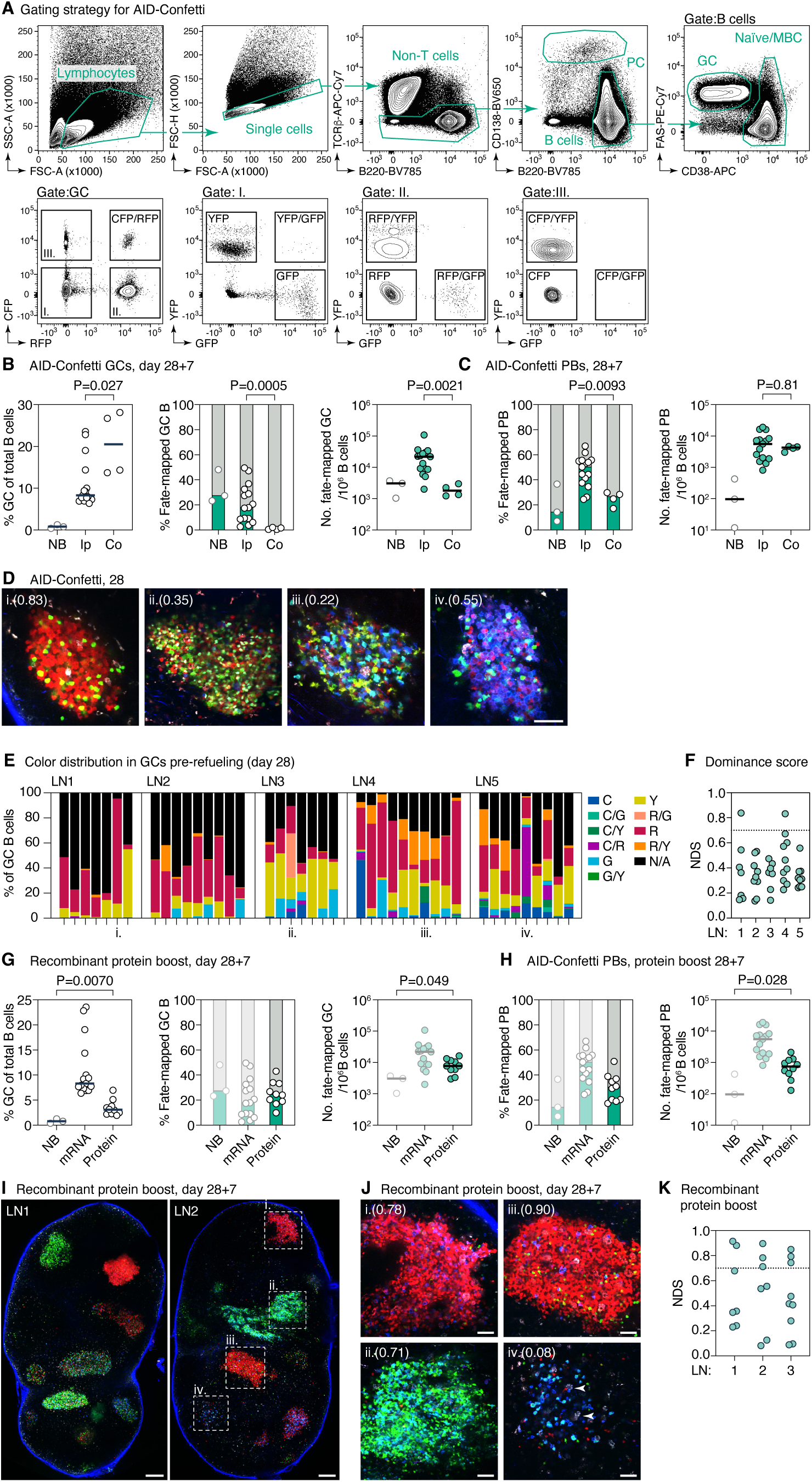
Tracking GC refueling using Aid-Confetti mice. **(A)** General gating strategy for AID-Confetti fate-mapping. **(B)** Quantification of flow cytometry of LN samples generated as in Fig. 1A, using AID-Confetti mice. Graphs show GC size (left), fraction (center), and number (right) of fate-mapped GC B cells. **(C)** As in (B), showing fraction (left) and number (right) of fate-mapped PB. NB, non-boosted; Ip, ipsilateral refueling; Co, contralateral boosting. Each symbol represents one mouse. P-values are for Mann-Whitney U test, only relevant comparisons are shown. **(D)** Representative multiphoton images of and single GCs from AID-Confetti mice at day 28 post-prime, prior to refueling. Details as in Fig. 2B. Scalebars, 50 µm. Numbers in parentheses are NDS. **(E)** Color distribution in individual GCs generated as in (D). Each bar represents one GC, colored by the fraction of cells expressing each color combination (color labels as in Fig. 2C). Roman numerals indicate GCs depicted in (D). **(F)** NDS for GCs in (D,E). Each symbol represents one GC. The dotted line represents a score of 0.7, above which a GC is regarded as a containing a clonal burst. **(G,H)** Quantification of flow cytometry as in (B,C), comparing mice primed with SARS-CoV-2 spike mRNA-LNP and refueled with either mRNA-LNP or recombinant spike protein. Data for NB and mRNA (lighter symbols) are reproduced from panels (B,C) for comparison. **(I,J)** Representative multiphoton images of cross-sections of whole LNs (I) and single GCs (J) from AID-Confetti mice at 7 days post-boosting with recombinant spike protein. Details as in Fig. 2A,B. Scalebars, 200 µm (I) and 50 µm (J). Numbers in parentheses are NDS. **(K)** NDS for GCs in (I,J). Each symbol represents one GC. The dotted line represents a score of 0.7, above which a GC is regarded as a containing a clonal burst.

**Figure S3.**
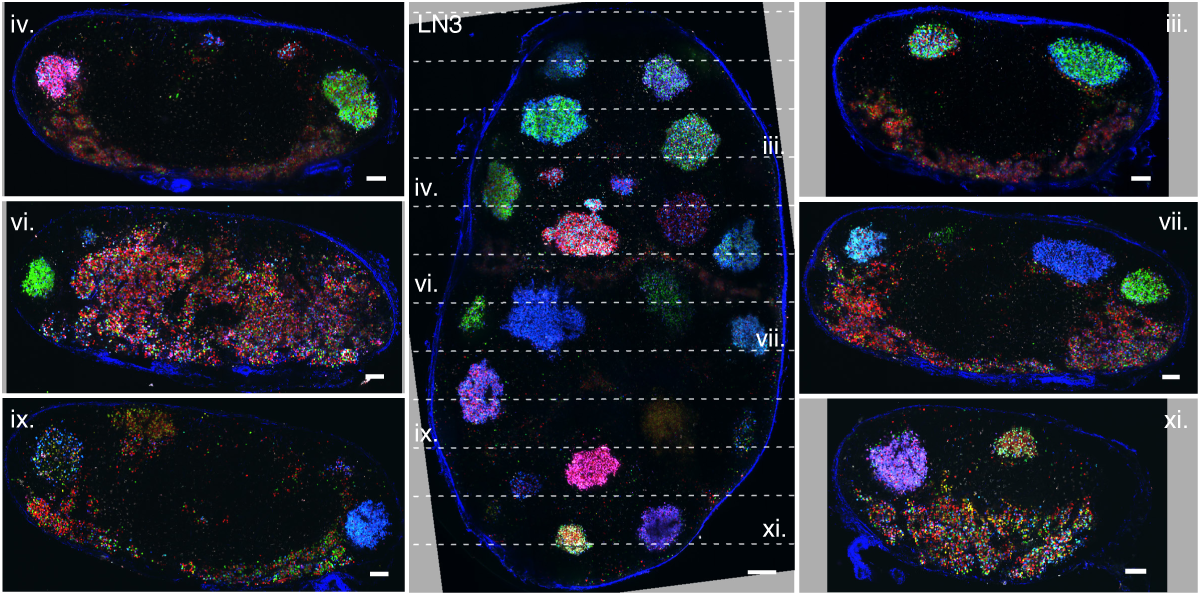
Multiphoton images exemplifying the process of LN sectioning for single-GC isolation. The central panel shows an intact GC (reproduced from Fig. 2A) prior to sectioning; panels on the right and left are LN slices imaged post-sectioning. Slices are numbered with roman numerals from the top. Scalebars, 200 µm (overview) and 100 µm (sections).

**Figure S4.**
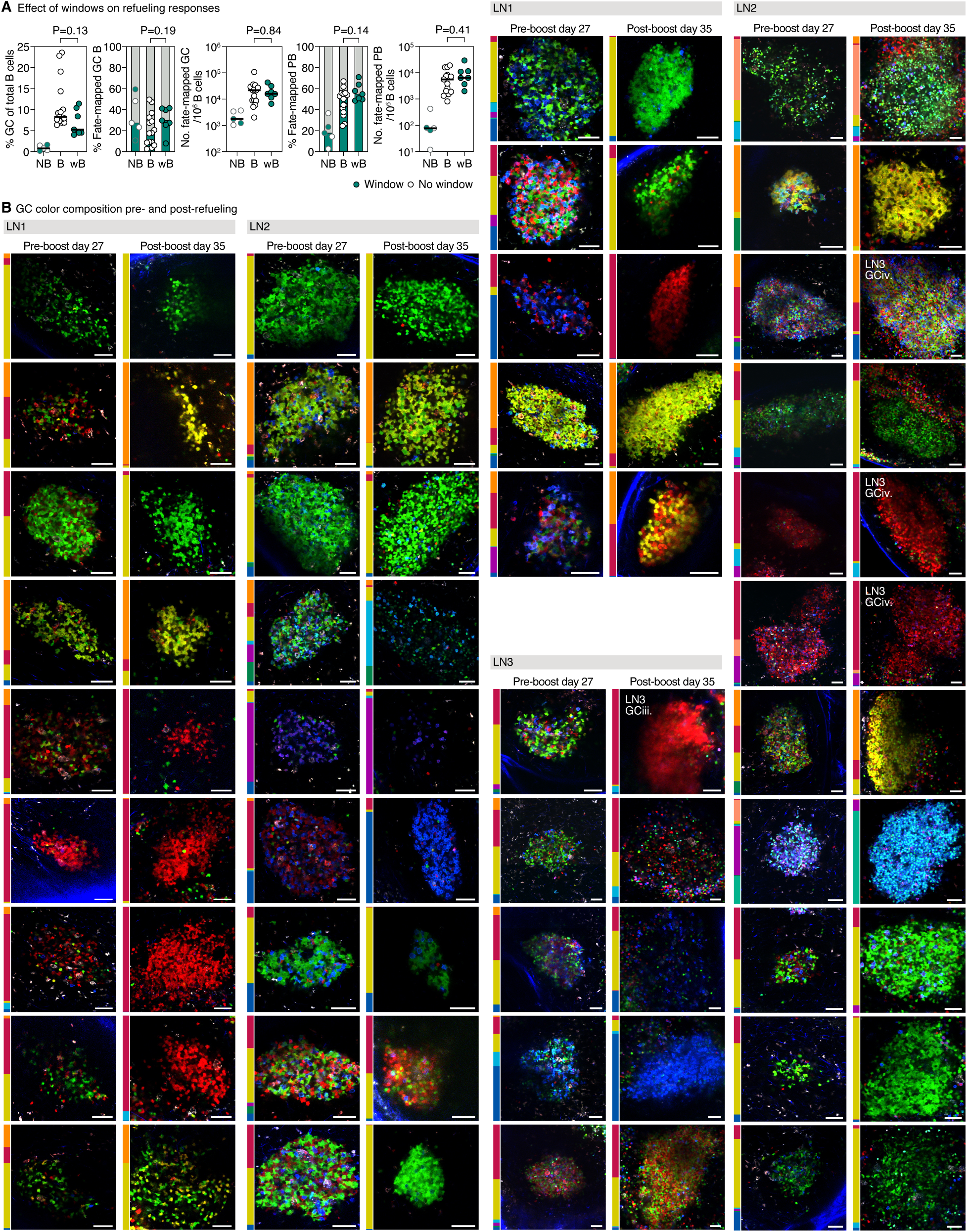
Imaging GCs sequentially with implanted inguinal LN windows. **(A)** Window implantation does not affect the response to refueling. Quantification of flow cytometry as in Fig. S2A,B, comparing the refueling response in mice with and without implanted windows. NB, non-boosted; B, boosted; wB, boosted following implantation and removal of imaging window as described in Fig. 3D. Each symbol represents one mouse. Dark symbols in NB are non-boosted GCs imaged 8 days following window implantation and removal. Data for non-window mice is reproduced from Fig. S2B for comparison and is shown in white symbols. **(B)** Full set of pre-/post-refueling pairs as in Fig. 3E (including those already shown in that image). Bars to the right of each image indicate the fraction of B cells of each fluorescent color in the adjacent image (color labels as in Fig. 2C). LN/GC names are indicated for GCs used for phylogenetic analysis in Fig. 4C.

**Figure S5.**
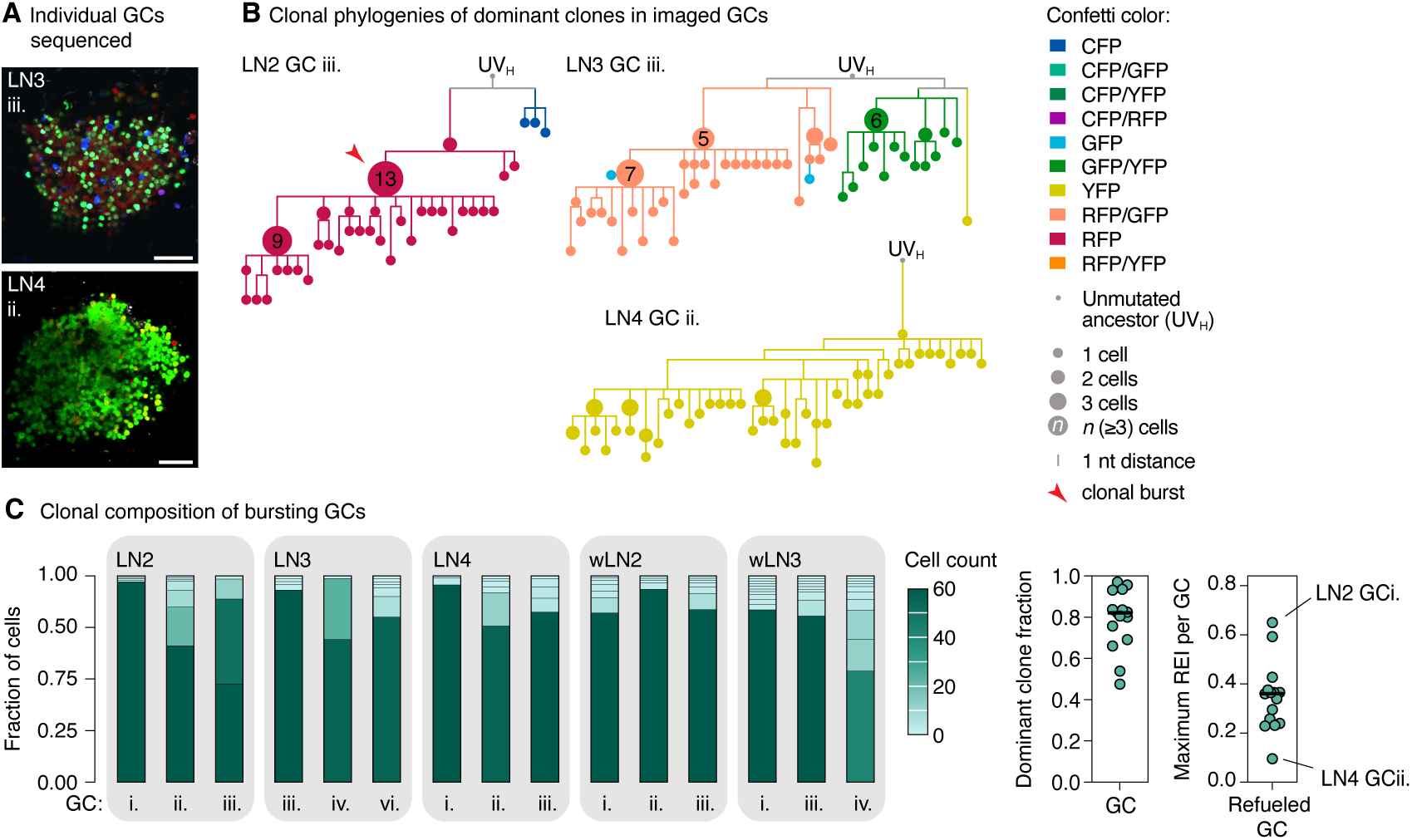
Additional phylogenetic trees from sequenced GCs. **(A)** Multiphoton images of GCs used to build phylogenies in (B) which were not previously represented in Figs. 2B and 4A. **(B)** Phylogenetic trees of the dominant B cell clones in the GCs depicted in panel (A), Fig. 2B, and Fig. 4A. Clonal bursts (defined as expanded nodes (> 1 cell) with REI > 0.25 [41]) are indicated by red arrowheads. **(C)** Clonal distribution of fate-mapped B cells in individual GCs. Each segment represents one clone, colored by the number of cells within the clone. The size of the dominant clone and the maximum REI in all 15 GCs are shown to the right, each symbol represents one GC.

**Figure S6.**
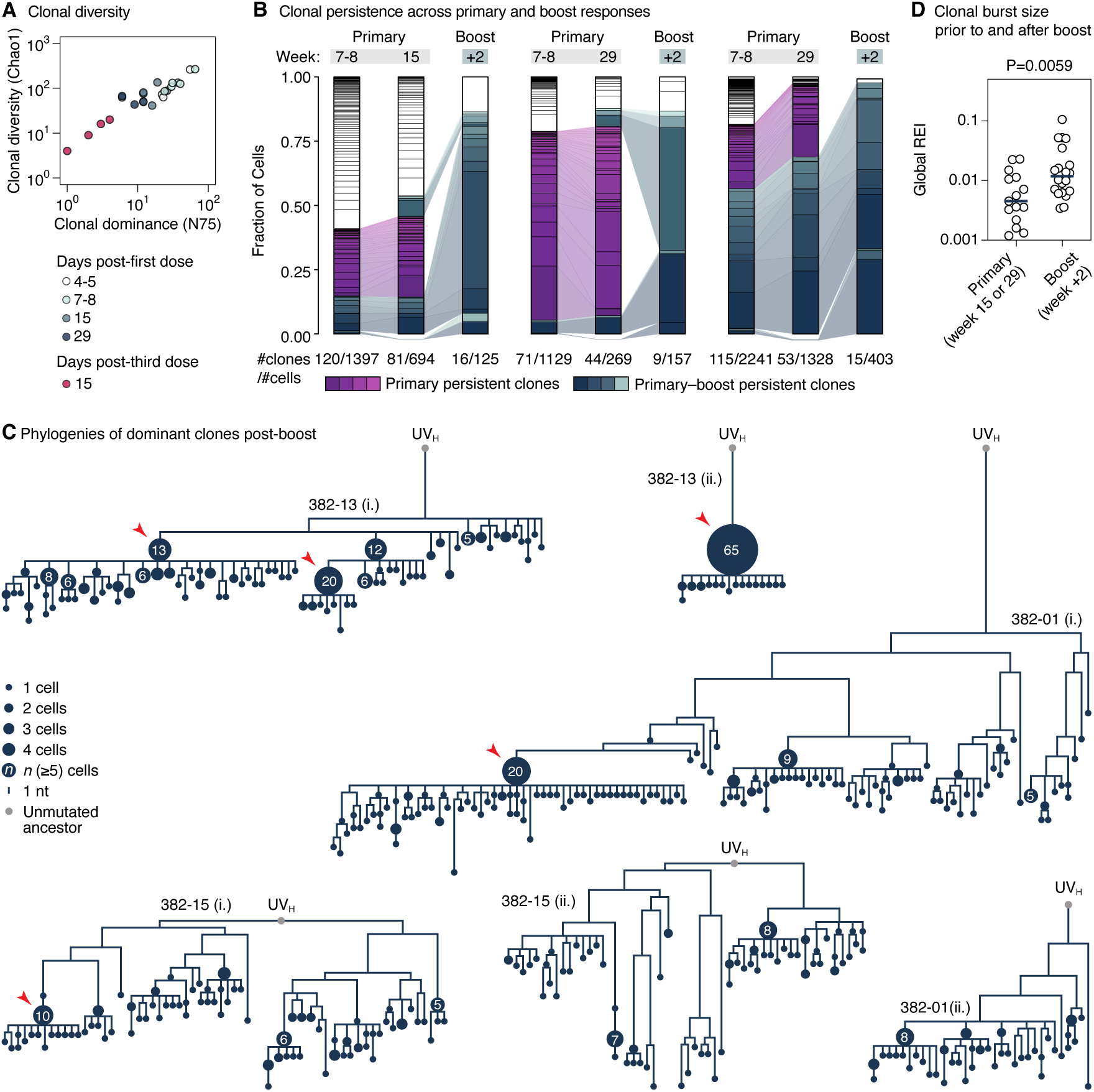
Clonal persistence and expansion in human post-refueling GCs. **(A)** Clonal diversity (Chao1 and N75) among spike-binding B cells. Includes data for all subjects in the dataset who had a post-boost FNA sample containing ≥ 50 GC B cells. **(B)** Clonality maps tracking the evolution of spike-binding GC B cells in human LN FNA samples at different time points prior to and after two primary-series (week 0, 3-4) and one booster (9-10 months) SARS-CoV-2 spike mRNA-LNP immunizations [25, 42]. Each segment represents one clone. Clones present in more than one compartment are connected and depicted in color; white clones were present in a single time point only. Numbers below each bar are (number of clones/number of cells) in each sample. Only subjects with two pre-boost time points and > 50 spike-binding cells in the post-boost sample were included, restricting the analysis to 3 subjects. **(C)** Clonal phylogenies for the 2 largest clones from each of the three subjects analyzed in (A), regardless of antigenic specificity. Arrowheads indicate burst points containing ≥ 10 cells in the parental node. **(D)** Global REI scores comparing clonal expansion sizes in pre- and post-refueling samples. Includes all clones containing ≥ 20 total cells and accounting for ≥ 2.0% of cells in their respective sample; samples as in (C). P-value is for Mann-Whitney U test.

**Figure S7.**
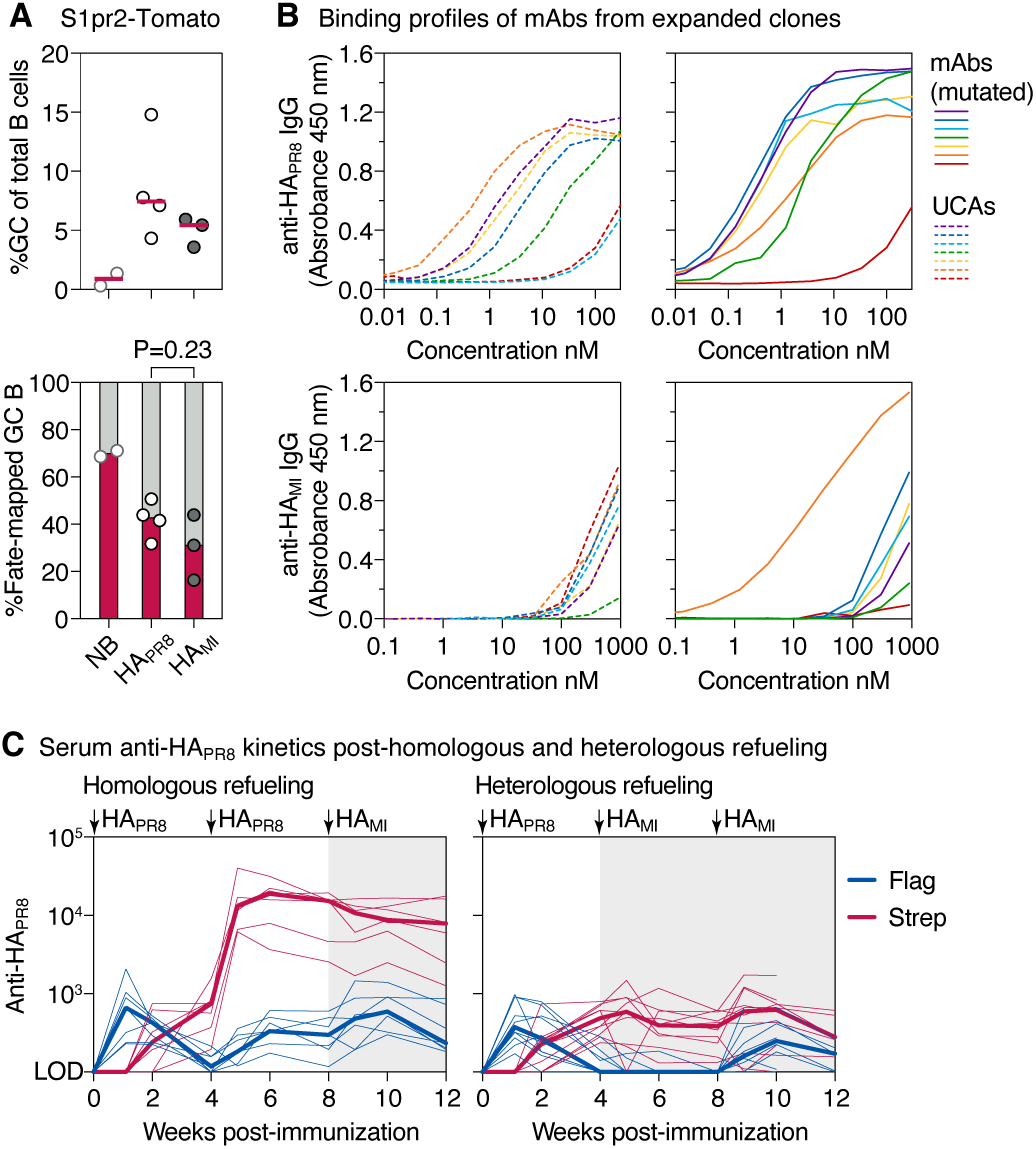
Heterologous refueling with an influenza hemagglutinin variant. **(A)** Quantification of flow cytometry (representative plots in Fig. S1B,C) of LN samples generated as in (Fig. 1A) using S1pr2-Tomato mice and upon homologous or heterologous HA refueling. GC size (top) and fraction of fate-mapped cells (bottom) are shown. Each symbol represents one mouse. P-values are for Mann-Whitney U test, only relevant comparisons are shown. **(B)** Binding to the indicated HA variant by ELISA of mAbs derived from expanded GC B cell clones sequenced from heterologously refueled mice. Summarized in Fig. 5D. **(C)** Tag-specific anti-HA_PR8_ serum titers in mice fate-mapped after the primary immunization (left) or at the time of refueling (right), measured by ELISA. Thin lines represent individual mice, thick lines link medians of log transformed titer values at each time point. Anti-HA_MI_ titers are shown in Fig. 5F.

## OTHER SUPPLEMENTAL MATERIAL

**Table S1.**
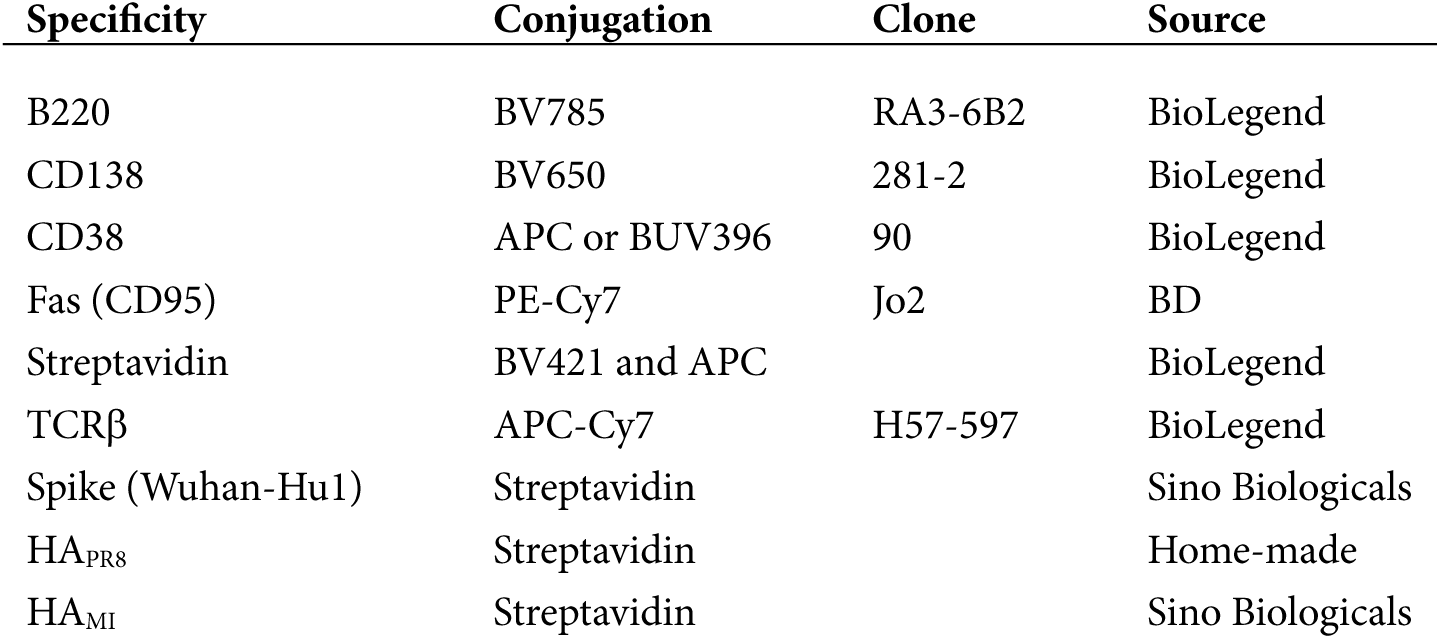
Reagents used for flow cytometry.

| Specificity | Conjugation | Clone | Source |
| --- | --- | --- | --- |
| B220 | BV785 | RA3-6B2 | BioLegend |
| CD138 | BV650 | 281-2 | BioLegend |
| CD38 | APC or BUV396 | 90 | BioLegend |
| Fas (CD95) | PE-Cy7 | Jo2 | BD |
| Streptavidin | BV421 and APC |  | BioLegend |
| TCRβ | APC-Cy7 | H57-597 | BioLegend |
| Spike (Wuhan-Hu1) | Streptavidin |  | Sino Biologicals |
| HA<sub>PR8</sub> | Streptavidin |  | Home-made |
| HA<sub>MI</sub> | Streptavidin |  | Sino Biologicals |

**Movie S1 (online only).** Z-dive through GCs from cortical to medullary. Scalebar, 50 µm; slices are 10 µm apart.

**Movie S2 (online only).** 3D rendering of sequentially imaged GCs showing pre- and post-refueling states. Scalebar, 50 µm.

**Spreadsheet S1 (online only).** Sequences of primers used for *Ig* sequencing.

**Spreadsheet S2 (online only).** Sequences of HA variants used for mRNA-LNP production.

